# The Role of Proton Transport in Gating Current in a Voltage Gated Ion Channel, as Shown by Quantum Calculations

**DOI:** 10.1101/371914

**Authors:** Alisher M Kariev, Michael E Green

## Abstract

Over two-thirds of a century ago, Hodgkin and Huxley proposed the existence of voltage gated ion channels (VGIC) to carry Na^+^ and K^+^ ions across the cell membrane to create the nerve impulse, in response to depolarization of the membrane. The channels have multiple physiological roles, and play a central role in a wide variety of diseases when they malfunction. The first channel structure was found by MacKinnon and coworkers in 1998. Subsequently the structure of a number of VGIC was determined in the open (ion conducting) state. This type of channel consists of four voltage sensing domains (VSD), each formed from four transmembrane (TM) segments, plus a pore domain through which ions move. Understanding the gating mechanism (how the channel opens and closes) requires structures. One TM segment (S4) has an arginine in every third position, with one such segment per domain. It is usually assumed that these arginines are all ionized, and in the resting state are held toward the intracellular side of the membrane by voltage across the membrane. They are assumed to move outward (extracellular direction) when released by depolarization of this voltage, producing a capacitive gating current and opening the channel. We suggest alternate interpretations of the evidence that led to these models. Measured gating current is the total charge displacement of all atoms in the VSD; we propose that the prime, but not sole, contributor is proton motion, not displacement of the charges on the arginines of S4. It is known that the VSD can conduct protons. Quantum calculations on the K_v_1.2 potassium channel VSD show how; the key is the amphoteric nature of the arginine side chain, which allows it to transfer a proton; this appears to be the first time the arginine side chain has had its amphoteric character considered. We have calculated one such proton transfer in detail: this proton starts from a tyrosine that can ionize, transferring to the NE of the third arginine on S4; that arginine’s NH then transfers a proton to a glutamate. The backbone remains static. A mutation predicted to affect the proton transfer has been qualitatively confirmed experimentally, from the change in the gating current-voltage curve. The total charge displacement in going from a normal closed potential of −70 mV across the membrane to 0 mV (open), is calculated to be approximately consistent with measured values, although the error limits on the calculation require caution in interpretation.

## INTRODUCTION

Hodgkin and Huxley(1-4) proposed that the nerve impulse consisted of the transport of Na^+^ and K^+^ across the nerve cell membrane, through protein channels that opened and closed in response to polarization or depolarization of the membrane voltage. Over the more than half-century since the channels have been discovered, the structure of the protein of which they are composed has been determined for the open state for several varieties of these channels(5-7), and a set of models has been proposed for the mechanism by which they open and close (the standard models referred to above); the common feature is the motion of one transmembrane (TM) helix, S4, orthogonal to the membrane, in response to depolarization of the membrane(8-14). This list of models is not exhaustive, but includes most of the most common versions. In addition to the gating mechanism, there are questions with regard to the transport mechanism of the ions through the pore, and as to the inactivation mechanism. Inactivation means the shutting down of ion transport without the channel returning to the fully closed position. There are both fast and slow inactivation mechanisms; the fast mechanism has been basically understood for some time(15), but the slow inactivation mechanism is still a subject of research. As we limit the discussion to activation gating, these topics will be discussed only to the extent that they interact with activation gating. The early work in the field has been comprehensively summarized by Hille(16). We will be mainly concerned with the voltage gated potassium (K_v_) channels, with some comparison with the proton conducting, but voltage gated, H_v_ channels. Both of these are found, in some form, in practically all organisms. Even viruses have been found to have a simple version(17). These channels exist in multiple forms, with some characteristics, such as the selectivity filters of potassium channels, conserved across practically all forms. There exists functional variation in channels found in different organs and organisms, but almost every cell must have potassium channels. The functional diversity and its possible implications has been discussed by Islas(18). Sodium channels, unlike potassium channels, are composed of a single peptide, and the four domains that comprise the channel are not identical in composition or function; these channels are sufficiently similar to potassium channels that results found on sodium channels are at least worth noting in a discussion of potassium channels as well, although it must be understood that they will not behave in quite the same manner. Studies of other potassium channels, like the bacterial KcsA channel, which is primarily pH gated and lacks a voltage sensing domain, are also informative. The structural similarities among these channels, especially the pore segments, is strong.

MacKinnon and coworkers produced a major advance in the field by providing the first structures of a channel, the KcsA channel just mentioned(5); the availability of structures has made it possible to discuss mechanisms at an atomic level. Since the KcsA channel, there have numerous structures determined. We will primarily discuss the 3Lut (pdb code) structure of the K_v_1.2 channel, for which the hydrogens were added by normal mode analysis(19). Fig. 1 shows the structure of the full channel, reproduced from the pdb website. Our calculations are so far limited to a large part of just one VSD. The calculations show the path for a proton through the section that has been calculated, with no water involved. However, once one gets past the hydrophobic section, there is water in the remainder of the path to the gate. This part has not yet been computed, but we can see what appears to be a clear pathway, and propose it as a hypothesis.

**Fig 1:**
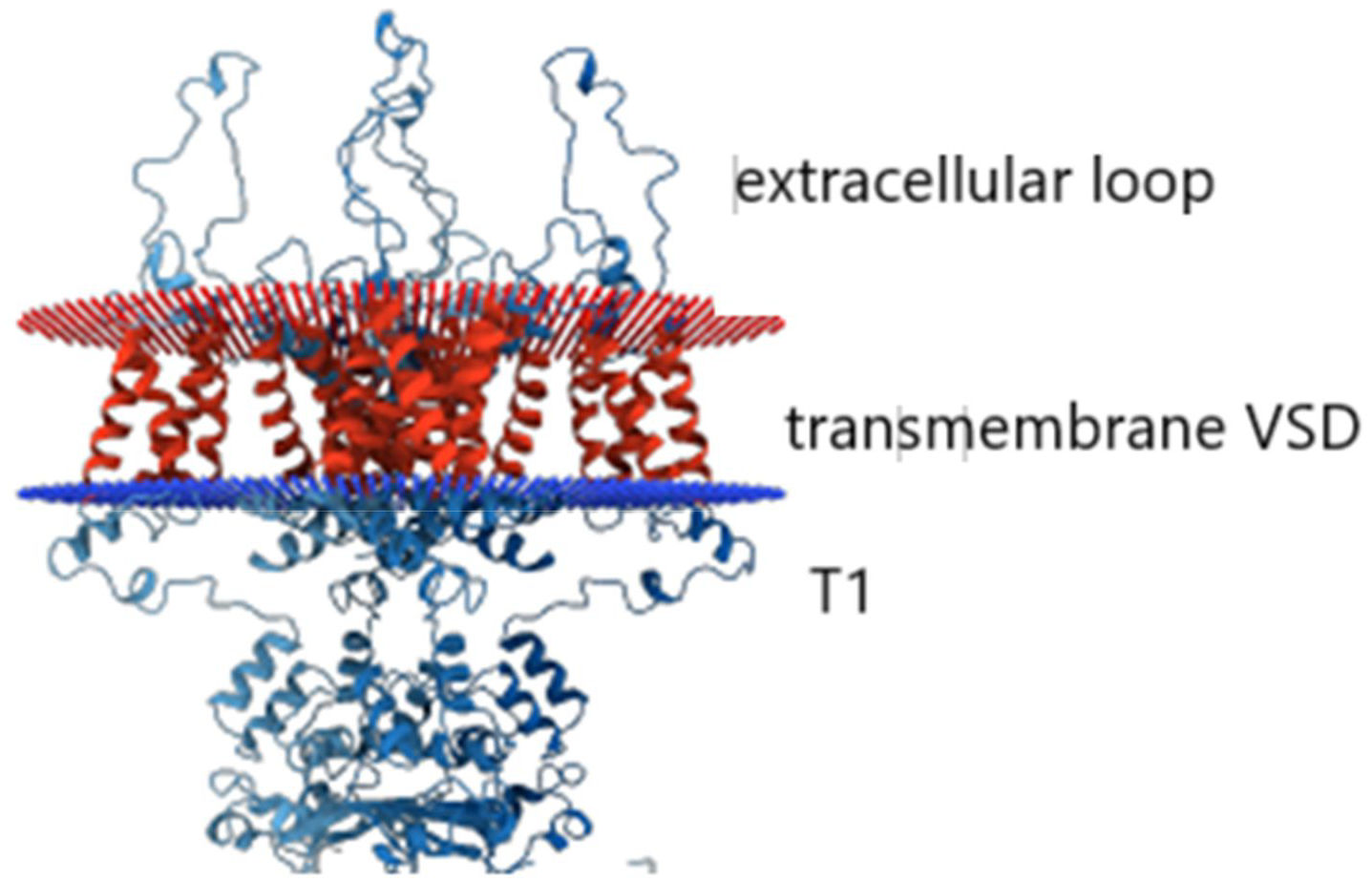
(reproduced from protein data base, rcsb code 3Lut(19): The red plane is the extracellular membrane surface, the blue plane the intracellular membrane surface. The extracellular loops connect TM segments, the voltage sensing domains (VSD, there are four, the one on the right labeled) each have four such segments, and the pore has 8 segments, two from each VSD. The T1 moiety below the intracellular membrane boundary is part of the postulated proton pathway; it is known experimentally to affect gating(22). The pore section is in the center.

Two quantities are fundamental in measurements on channel gating: gating current and gating charge. Prior to opening the channel, there is a capacitative current of relatively short duration, the gating current(20). This is consistent with the motion of positive charges through the VSD of the channel; the potassium channels we are considering have four such domains, plus a central pore domain through which the ionic current passes. The gating current passes through the VSD; the gating charge can be inferred from measurement of the gating current; this is often done by assuming a single “Boltzmann curve”:

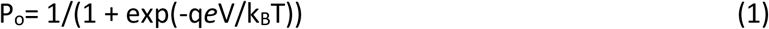

where P_o_ is the open probability for the channel, q is the gating charge, *e* the electronic charge, V the applied voltage, k_B_ the Boltzmann constant, and T the temperature. This produces a saturating curve with increasing voltage, which is followed quite closely by channels, and fitting the curve can give q. However, assuming a single step for gating is almost certainly incorrect, and using this assumption to determine gating current may be a mistake (21).

Each VSD has four TM helical segments; segment S4, which connects through a linker to the pore domain, has 4 arginines (the number is not 4 in all channels, but, if S4 motion were responsible for gating charges, generally about three would have to cross the membrane field to produce the measured gating charge). The arginines are assumed to be positively charged (the solution pK_a_ is close to 12), so taking the motion of these charges to constitute the gating current was a natural assumption. Our proposed gating mechanism does not fit in the group of standard models, since it does not have the S4 TM helix providing the gating current by a large scale movement with its putatively positive arginines. We show that the arginines need not always be charged in the membrane context; a proton from the arginine can transfer to a neighboring glutamate. This is one step in a proton current that would be detected as gating current, although the total charge displacement is actually the gating current, and is not limited to movement of specific charged entities. No such proton transfer is contemplated in any standard model. We provide quantum calculations as part of the evidence to support this proton transfer. The VSDs for the various channels that belong to this family are not all alike. The H_v_1 channel is, in its entirety, a close analog of the VSD; it is a proton channel. Its structure, to a reasonable approximation, is that of a dimer of a VSD of a potassium channel, especially at the extracellular end. Some other channels appear to have proton transmission sections that are also analogous, although they are not as closely related as H_v_1; some are listed in a later section.

The evidence that gave rise to the standard models in which positively charged S4 moves to produce gating current can be reinterpreted so as to be consistent with proton motion. The primary evidence came from experiments in which the arginines on S4 that were believed to produce the charge that moved were one by one mutated to cysteine. The cysteine side chain, if ionized, consisted of one negatively charged sulfur atom that could react with methanethiosulfonate (MTS) reagents; the unionized –SH side chain could not react, and this fact is crucial to interpretation of the results. Added MTS reagents were observed in some cases to kill the channel, while in other cases they failed to do so. This “scanning cysteine accessibility method” (SCAM)(23, 24), includes an assumption that is key to the interpretation: the dielectric constant inside the protein was assumed to be too low for the cysteine to ionize, so the reaction with MTS (which must have taken place to kill the channel) was assumed to occur at the aqueous interface of the membrane, meaning that S4 must have moved enough for the cys to reach the surface to react. It was found that the reactive arginines appeared to shift with the voltage in approximately the way that might have been expected if the S4 had moved. If the MTS reagent were applied from the extracellular side, more cysteines reacted when the voltage was off, while intracellular MTS was more likely to react at negative voltage, when S4 should be at the intracellular side to close the channel. Thus, the S4 seemingly moved to the extracellular surface when the voltage was removed, and to the intracellular surface when the voltage was applied, a method first used on channels by Yang and Horn (25). The method, or variants on it, have been used by multiple workers since the Yang and Horn paper. However, before this interpretation can be accepted, we note that the side chain of arginine is more than twice the volume of the full cysteine side chain(26) (see Fig. 2), and this is not counting the decrease in volume of cysteine upon ionization. The entire cysteine side chain is an S and an H, so ionization leaving only S makes a significant difference. The difference in volume from arginine (depending somewhat on the definition of the volume) is roughly 75 Ȧ^3^. This is enough for the head group of the MTS reagent, if the cysteine does not need to go to the surface to ionize. This difference is also about the volume of 2-3 water molecules. If some water enters the cavity which is left by the mutation, ionization can occur *in situ,* so that a central assumption of the method is not necessarily supported in this system; local changes in side chain rotation would be enough to account for the results. Thus the R➔C mutation results are not definitive; they may be interpreted in a manner different from what has been assumed. There may be another problem: the protein structure may undergo a local collapse into the cavity created by the mutation to cysteine (this might happen with some other mutations to cysteine as well). Thus the interpretation is ambiguous in more than one way.

**Fig. 2:**
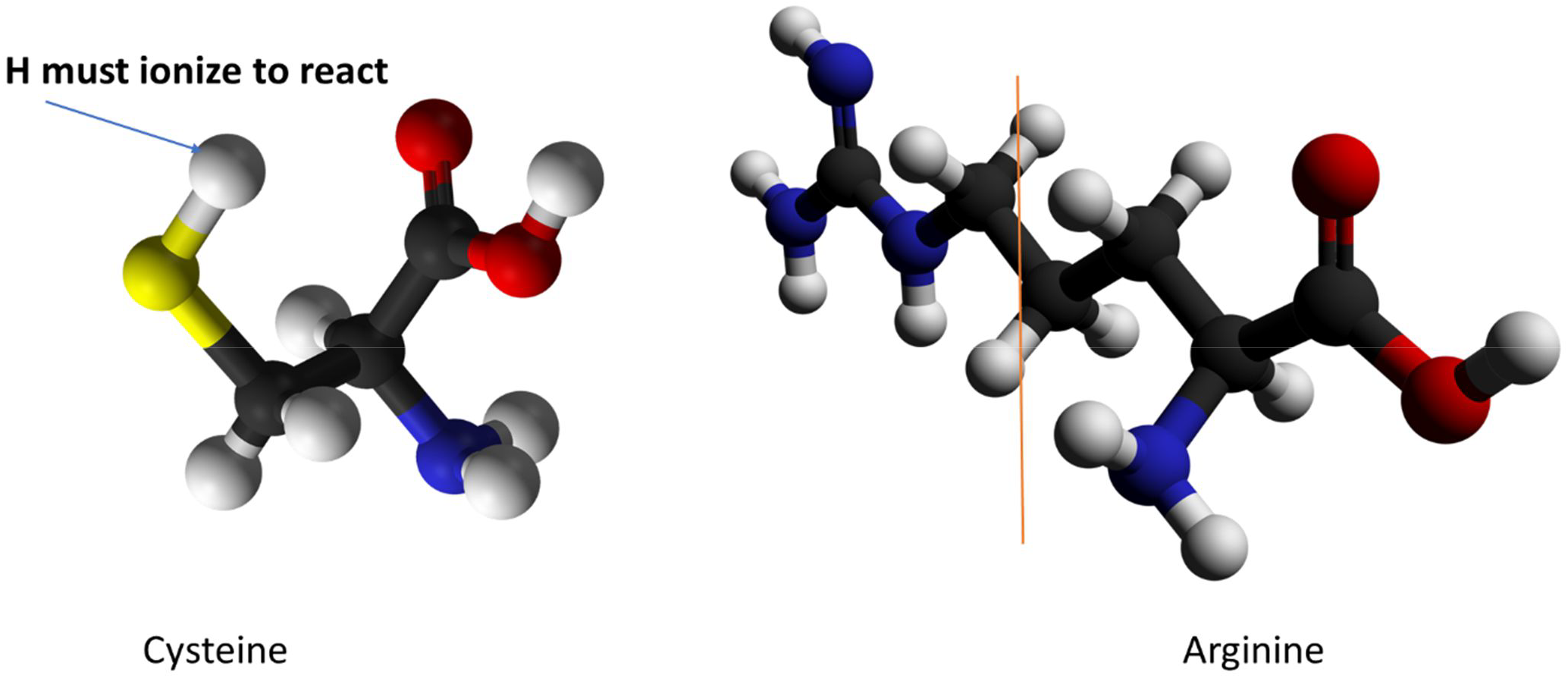
Comparison of cysteine and arginine: the section to the right of the vertical brown line on arginine approximates the volume that would accommodate a cysteine. If the hydrogen on the cysteine –SH ionizes, the cysteine becomes even smaller. The entire guanidinium group plus the proximate carbon take up space greater than the cysteine, and adequate for 2 to 3 water molecules.

A second line of evidence that has been interpreted in terms of the standard models comes from fluorescence quenching. FRET (fluorescence resonance energy transfer) or LRET (lanthanide resonance energy transfer—often Tb^3+^ is used as the lanthanide) measurements give an indication of the distance between residues, which appears to change with applied voltage (27-34). Again, there are difficulties with this type of measurement. These have been discussed in the literature, and major efforts to overcome these difficulties have been made. However, side chain rotations, loss of salt bridges, and, for that matter, proton transfers, would affect the results, so that the error limits on these measurements must be considered to be comparable to the distances measured. As these errors are well known, we will not discuss them. LRET is somewhat better than standard FRET, but does not overcome the local structural effects of mutation. Undoubtedly there is value to these measurements, but they do not necessarily lead to high resolution results. The measurements are made on a mutant channel, which may be a few Angstroms different in spacing from the wild type, a problem for a measurement of distances in which 3 Å is a major difference. This said there is evidence from these experiments that does not necessarily fit with the standard models. Posson et al found a vertical movement of only 2 Å for S4(35); they proposed a model in which the electric field was refocused in such a way as to produce a gating charge of 13*e*, in accord with the measured value for the *Shaker* channel. Part of the reason this could be done was that it was shown, by use of a solvatochromic dye, that the field was very large at the location of a single arginine residue(36). Related data showing that the motion of S4 was very limited was added by a series of LRET measurements by Chanda et al(32). Slightly earlier work by Cha et al also led to the same conclusion(37). There are additional problems with assigning gating current, as its exact value is something of a problem. Ishida et al measured the gating current for a channel that was supposed to be very similar, K_v_1.2, and found a gating charge of only 10*e* (38); based on mutation experiments, they concluded that the gating current was produced by a different mechanism from that of *Shaker*. There are multiple states through which the channel passes on its way to opening(21); essentially all models require multiple states, and this is consistent with the measured kinetics, although not all results lead to exactly the same number of states

In addition to experiments, molecular dynamics (MD) calculations have been used as further evidence for the standard models. MD calculations use classical force fields; these force fields include electrostatic terms, as well as kinetic energy, and empirical force terms that should, in principle, account for all other energy terms (van der Waals, torsion, etc.) However, quantum calculations seem to be required because the exchange and correlation energy terms, which have no classical analogues, have differences between similar positions of protons or side chains of the order of 10k_B_T, so cannot be ignored. Proton shifts are not possible with MD, nor is charge transfer. Seemingly symmetric atoms also do not have the charge symmetry that is typically assumed, e.g., the amines (NH1 and NH2 in the pdb notation) that are part of the guanidinium ion; these effects are small on a percentage basis, but, leading to energy differences larger than k_B_T, would lead to differences in structure that cannot be taken into account in a strictly classical computation. Even including Drude polarizability(39) does not cure this problem. There is no way to compare the classical energy to the QM energy, as the Coulomb and the kinetic energy terms are computed on a different basis. There is further discussion below, in the section on Computational Methods, where the differences between classical and quantum calculations are also discussed.

Can the evidence that has been accumulated over the past several decades be interpreted other than by the standard model? We have offered an alternative, proton transport, which would provide the gating current, and would be able to open and close the gate (40-45), but must show that the evidence that has been interpreted in terms of the standard model can be interpreted without it.

## EVIDENCE THAT CAN BE INTERPRETED WITHOUT INVOKING S4 MOTION

It is necessary to summarize this before we go further:

1. There is considerable evidence that protons can go through the VSD, as well as through the closely analogous H_v_1 channel, and other channels discussed below. H_v_1 is a proton channel, but the extracellular section, more than half the channel, is very similar to the VSD (46-51). The intracellular part of the K_v_1.2 VSD differs from the corresponding section of the H_v_1 channel, as the path that the protons must follow is different in the two cases for that section of the channel. In K_v_1 the proton must continue on to the gate of the channel, moving parallel to the membrane, while in H_v_1 the proton goes directly through the membrane to be removed from the cell. However, the similarity of the two types of channels is a strong indication that the structure of the VSD is also suitable for transmitting protons. Mony et al used fluorescence methods to suggest a role for the S1 TM helix in H_v_1 proton transport, and thus in forming a proton channel in K_v_ channels, by analogy; they interpreted the proton motion as being independent of gating(52); Goldshen-Ohm and Chanda noted this as an interesting result, but pointed out that the evidence was not conclusive (53) (they did not suggest any connection between proton current and gating, rather that the role of S1 was not conclusive). Further evidence comes from mutation of the end arginines, especially the most extracellular one, to histidine (54-56). In this case, the VSD becomes a proton channel; therefore, the end arginine seems to have the function of preventing the protons that would be moved internally from exiting the VSD; by limiting the proton trajectory, these arginines would leave the protons to produce a capacitative current in response to application of a field, which would produce the same phenomenological result as S4 motion is considered to in the standard models. It has also been shown that mutating the amino acid next to the arginine to histidine does *not* produce a proton current; the proton is not simply carried across the end of the segment by the histidine moving up or down; if the histidine moved, one would expect such a mutation to provide a current comparable to that of the arginine mutation. The fact that it does not supports the idea that S4 does not move, but the protons do. Much more recently, Zhao and Blunck(57) have shown that the VSD can transport protons (and, to a limited extent, some other cations). Therefore, the idea that protons move through the VSD is not controversial, although the idea that this has to do with gating is. How it is possible to account for the motion of protons in response to voltage while having additional charges move to provide the gating current, in standard models, is not entirely clear. Somehow, the protons would have to not contribute to gating current even when they move.
2. In a related channel which a K^+^ channel controls a phosphatase enzyme, belonging to the parasite *Ciona intestinalis*, the Ci-VSP channel, R➔H mutations allowed proton transport completely through the section closely analogous to the VSD of K channels. This appears similar to the *Shaker* case, where Starace and Bezanilla used an R➔H mutation(58) to produce a proton channel from the VSD, the case just discussed. Villalba-Galea et al have discussed the Ci-VSP channel in some detail (59), as have Sakata et al(60). If two residues interact, this provides important information as to mechanism. One method of determining whether there is interaction or not is to put spin labels on two locations where interactions are thought to occur, one can find whether the electron paramagnetic resonance (EPR) spectrum of one label is perturbed by another, which implies that the two labels are relatively close to each other. This may change with gating, suggesting that there may have been motion by the labels. The technique is subject to the usual problems of having a large foreign residue inside the VSD. Displacement of the native residues by the probe, as the probe is large, is a consideration. The difference in the way the probe rotates as the channel gates is another. These are problems similar to FRET, with its large probe. The technique was described by Perozo et al in 2002 (61, 62). There have been a number of applications, especially, but not exclusively, to ligand gated channels(63, 64). However, Li et al studied the VSD associated with the Ci-VSP, using both X-ray and EPR(65); here the VSD is fairly much like that of the potassium channels. This channel is almost unique among K^+^ channels in that much greater depolarization, actually inverse polarization, is required to open them. They are still closed at 0 mV, which makes it possible to get a closed conformation without applying a voltage. These authors concluded, by comparing the structures at 0 mV (closed) and +100mV (open) that the motion of S4 was about 5 Å, which they associated with a “one-click” translation (meaning an arginine moved to the position of the next salt bridge to close the channel). EPR gave both dynamic and structural information. Of course, the spin probe that was attached was larger than 5 Å, so a smaller motion is plausible, as is a larger motion. This channel may also not be the same as the standard channels, as can be told from the fact that it is closed at 0 mV, and some of the studies are on an R➔E mutant. Research is continuing on this channel.
3. There are other proteins that have some sections of their internal structures very similar to those of the VSD, and that are known to transmit protons, including cytochrome c(66-68), bacteriorhodopsin(69-71), and the M2 channel of the flu virus(72, 73). Of the latter cases, some have water, unlike the VSD amino acid triad that can transmit a proton without water.
4. Nguyen and Horn(74) showed that S3, which has negative charges, appears to move in the same direction as S4 when the standard cysteine mutation plus MTS experiment is done; this would indicate that this experiment measures *in situ* availability of the residues, but the authors found an explanation consistent with the standard model. The possible motion looks a little like the MacKinnon paddle model in which half of S3 moves together with S4(75, 76); a number of objections have been raised to this model, and it would take too long to discuss them all here; this model at present does not seem to be a probable candidate for gating.
5. Naranjo and coworkers found that making a mutation of either sign(77) appears to have the same effect on gating charge; if S4 moves outward to open the channel, a positive (base) mutation should add to gating charge, while an acid, if ionized, hence negative, should subtract, but both subtracted. If S4 motion has nothing to do with gating current, this is easily understandable, but the authors found an interpretation that appeared consistent with the standard models; it is less clear that the interpretation was consistent with other experiments in which S4 would have to make somewhat different motions.
6. It has been shown that the first step in gating is a very small and very fast component of gating current (about 1% of gating current, and faster than the RC time constant of the membrane, hence too fast to measure, <2μs rise time), the “piquito”(78, 79). For a standard model interpretation one must assume that it is some sort of side chain rearrangement in an energy landscape; however, the nature of this energy landscape remains to be fully defined (80). An alternate interpretation is as a first step in a proton cascade, in which a proton tunnels to a neighboring residue, with the voltage change making this possible by matching energy levels. Such a component of gating current makes little sense in the standard models. However, Green and coworkers(81, 82) found that this could be explained if there is a threshold for the first transition leading to the start of gating current; the standard model seems to rule out thresholds because the current-voltage gating current curve fits a Boltzmann curve for a single transition (as previously noted, this is an oversimplification (21): see eqn. 1). However, if the threshold is distributed as an nearly Gaussian curve of width approximately k_B_T, then one gets a very similar curve(83). This said, the piquito, with threshold, does not rule out the standard models, if the rise time is not too much faster than 1 μs.
7. Fohlmeister and Adelman found what appeared to be a higher harmonic response to a large sine wave imposed on the membrane voltage, in a membrane with multiple sodium channels(84, 85). This experiment has been largely ignored, but unless there is some problem with the experiment itself, for which there is no evidence at this point, it deserves to be taken seriously. The threshold discussed by Green, however, turns out to be consistent with this experiment(83).
8. In addition to these results, some LRET and FRET experiments discussed previously suggest a small vertical movement, smaller than the standard model seems to require (32, 35, 37). Although not all such experiments lead to small apparent movement, the fact that some do suggests using care in interpreting all the results. Small movements would allow the gating current to exist if the field dropped across only a very small distance (although even then some care in setting up the model is required).(36)
9. In a standard model, at least in some versions, the first arginine does not contribute to gating current. Presumably this arginine would not move. This seems entirely reasonable, as it reaches into the headgroup section of the membrane. Arginine can complex with phosphate, and the headgroup region has a phosphate group for every lipid; not all lipids have net charge, although some negatively charged lipids are needed(86, 87). Furthermore, phosphatidylinositol 4,5-bisphosphate (PIP_2_) is needed to have a functioning channel(88-90), and this adds negative charges. While evidence that PIP2 is involved in complexing arginine in channels has not been reported (yet), this would be consistent with the existence of phosphate-arginine complexes; so far a complete proof awaits further evidence. However, calculations(91, 92) and experiment(93, 94) both show that guanidinium and phosphate form complexes that are fairly stable. Therefore, the end arginine, which reaches into the headgroup region, would be complexed with phosphate, with high probability. It would be necessary to release this complex if the S4 were to move as a rigid body, or nearly so. With the first arginine anchored in the membrane, there remains the possibility that S4 could break. With a fraction moving down, while the extracellular section remains in place, the broken helix would have to re-form thousands of times with no mistakes at all, such as returning to a local minimum with a different set of hydrogen bonds and salt bridges; returning so many times without error seems very implausible. The reason the S4 helix would have to re-form without error thousands of times can be understood as follows: consider that the channel must open and close more than once per second in many nerve cells, some of which fire at rates of 50 Hz. Let us assume 10 Hz to be conservative. A minimum lifetime for the channel might be 10 minutes; even this is expensive, in terms of the energy the cell must expend, if the protein must be resynthesized that often, but let us say 10 minutes. Then the channel must open and close completely accurately 6000 times in its lifetime, and with four transmembrane domains, that means a total of 24,000 openings and closings before the protein is replaced. If the S4 really must re-fold exactly correctly that many times, it must somehow use the rest of the VSD as a chaperonin, something for which there is no evidence, nor does it seem clear how it could be possible. This supports the idea that S4 does not unfold. However, if it does not, then there must be some way for the first arginine to get out of the headgroup layer, something that also seems extremely unlikely.

Besides these specific experimental results, there is additional experimental evidence that merits consideration.

## TEMPERATURE DEPENDENCE OF GATING

Temperature is one of the most obvious variables to test; it sets limits on the possible range of activation energy. If we have a single step to gate, it would determine the activation energy. The temperature dependence also shows when we are likely to have more than one step. In this case, there is good evidence that there are multiple steps. We can list here only a handful of the multiple studies that have varied temperature. Q_10_ is defined for gating as the change in open probability over 10°C, and for conductivity the change in conductivity over 10°C; for an open probability that follows Arrhenius behavior this can be translated into an activation energy. For potassium channels conductivity Q_10_ values in the range of 1.5 to 1.9 are found(95). For a proton channel, the values vary from 2.2 (10 °C) to 1.3 (40°C), suggesting the existence of different rate limiting steps for transport at low and high temperature(96) (activation energy of roughly 50 kJ (low temperature) vs 20 kJ (high temperature. A Q_10_ of 1.7 from 290K to 300K corresponds to 40kJ, or about 16 k_B_T. Temperature dependence has been studied by other methods, and with varying results. Rodriguez et al showed that there are several different regimes with varying amounts of charge transmitted, and Q_10_ values that were <2 in some regions, and ≈3 at higher temperatures(97). Gating has a larger Q_10_. Chowdhury and Chanda found several steps with Q_10_ values for gating current that gave an overall activation free energy of 58 kJ mol^-1^(98). Different steps have different Q_10_ values (hence, different activation energy). The temperature dependence of gating is larger than that for conduction. Rodriguez et al found an activation energy of about 100-110 kJ(99) for gating.

Some channels show much greater deviations from Arrhenius behavior; for example the TRP channels, which sense temperature, have Q_10_ over 30 in the range in which they sense temperature(100). However, for the type of channels we are discussing here, the value of Q_10_ in the physiological temperature range tends to be a little less than 2 for conduction, and approximately 4 for open probability. For the temperature sensing TRP channels. the mechanism of gating must be different; it appears so steep a temperature dependence requires a cooperative mechanism (but see(100) for a proposed mechanism). There is no evidence that gating in the non-TRP channels is cooperative, and no direct evidence that pertains to TRP channels either.

Fleming, Moon, and coworkers have considered the thermodynamics of inserting various side chains into a membrane(101). Their focus has been on bacterial outer membrane proteins (OmpA, and other outer membrane proteins, which are essentially beta barrels that allow substances through the outer membrane of gram negative bacteria). The relevance to the K_v_ VSD is that they find that it is fairly difficult to insert arginine, considered as a charged residue. If gating required charged residues to move through the headgroup region into the extracellular space to open the channel, or else intracellularly to close it, it would be difficult. While these authors considered only the insertion of the protein (unidirectional) to the membrane, the channel, if it leaves and reenters would not be stabilized by a drop in free energy, since it must go up the free energy gradient in one direction. This suggests that the extent of travel of S4, on the standard assumption that the arginines are charged, must be limited to keeping the charges within the membrane, and almost certainly within the hydrophobic part of the membrane. If the arginines are not charged this does not apply, but then the arginines would not contribute to gating current.

## K^+^ CONCENTRATION DEPENDENCE OF CONDUCTION

The dependence of the conductance of a KcsA channel on the concentration of K^+^ ion is similar to that of other K^+^ channels. It was measured by LeMasurier et al(102); the data from that paper has been replotted by Kariev and Green as Fig. 3, using −RT log [K^+^] = ∆G(40). This shows the conductivity is proportional to the free energy of the [K^+^] ion, which is consistent with the existence of a single barrier for conduction of the ion. The free energy is given relative to the low concentration, hence is shown as ∆∆G. The conductivity is σ, and is shown as relative conductance. This would be consistent with a barrier at the entrance to the pore, if the conductivity measures the effective concentration of the K^+^ in the pore, and the free energy is proportional, as just noted, to the ln[K^+^]. In considering the thermodynamics of the channel, this contribution must be included. The existence of a barrier at the pore for an incoming ion has implications for gating as well. If a gate becomes too wide open, such a barrier is hard to understand; if there is only a limited extent of motion at the gate, the existence of such a barrier is much easier to understand.

**Fig. 3:**
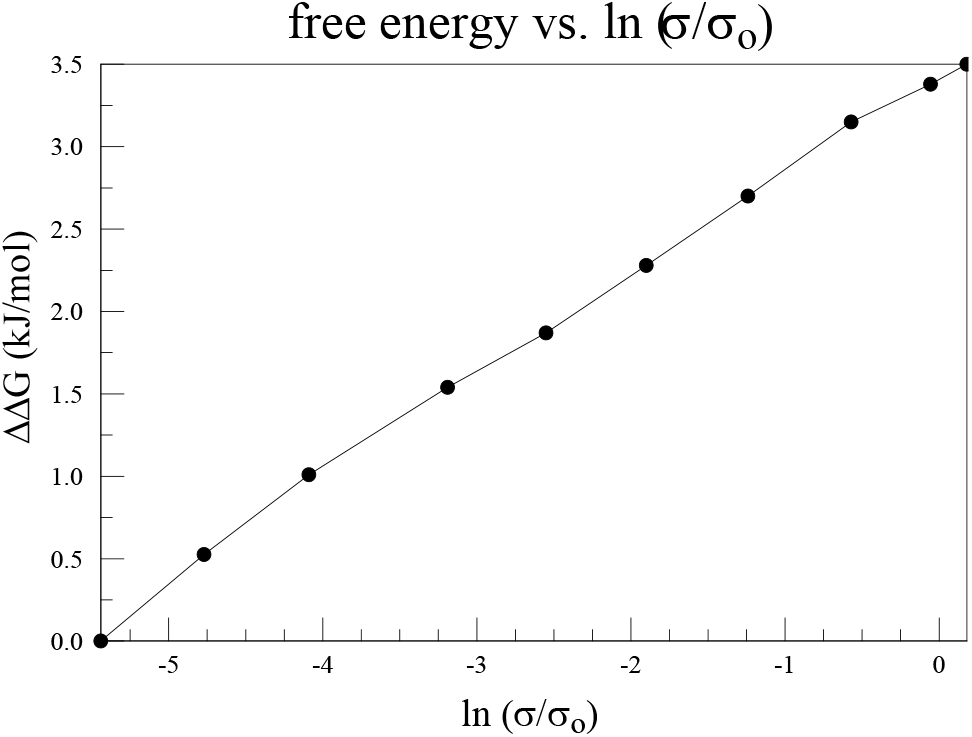
The conductivity of the KcsA channel, like other potassium channels, is proportional to the log[K^+^], hence to the free energy of the potassium ion in solution. Reprinted from Kariev and Green (40).

## THERMODYNAMICS OF AMINO ACID INTERACTIONS

it should be possible to find whether two residues interact by looking at the change in open probability (P_o_), and gating current and charge, with mutations of the residues, through mutant cycle analysis. One can also determine the free energy changes associated with the mutations. In this method one performs two single mutations, and then the double mutation. If the double mutation ∆G is not equal to the sum of the two single mutation ∆G values, one concludes that the two residues must interact. Here, the cycle would include states as shown by Fig. 4 (except for notation, this is essentially the same diagram as is given in many papers on mutant cycle analysis). A brief summary of this type of analysis, under the name “linkage analysis”, by Shem-Ad and Yifrach, provides a useful discussion of the method (103); these authors trace its conceptual history back to Wyman(104), in 1964.

**Fig. 4:**
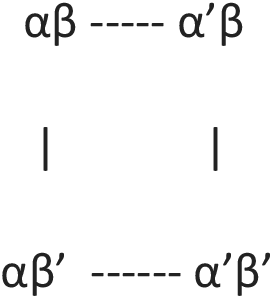
Schematic diagram for mutant cycle (linkage) analysis

The ∆G values around the cycle must add to zero, so that mutating the α and β residues to α’ and β’ allows one to determine the interaction free energy, as estimated from the measured variables, in open and closed states. Second, it allows an estimate of changes when the mutated residues are used, first separately, then together. The free energy change in one component (α) is ∆G_1_ if component β is present, ∆G_2_ if β’ is. Let ∆∆G be the difference ∆G_1_ – ∆G_2_. If ∆∆G is appreciable, presumably the residues interact. Otherwise, (∆∆G small or zero) they do not. Shem-Ad and Yifrach apply this to open and closed states of a generic channel, while Sadovsky and Yifrach (105) give a specific example involving a third residue (the diagram above then becomes a cube). Several examples have been considered by Chowdhury and Chanda, and they have reviewed the subject(106). The same authors have suggested a method of estimating free energy using the median voltage for activation(107). Because this subject has been reviewed, we will limit our discussion. It is not possible to review all the work done with double mutant cycle analysis on K_v_ and H_v_ channels, but a fairly representative sample can be cited (108-116); several of these papers consider the hERG potassium channel, which is found in heart in humans. Papers on Na_v_ channels in this context are just different enough that they would require a separate discussion, and there are fewer such papers. The question of which residues to study requires a model of which residues are of interest, as it is obviously not possible to study all possible pairs in a channel. Sadovsky and YIfrach (105), considered a three residue interaction. In addition, there is the possibility that the pair of residues studied may affect other residues so as to alter the local environment, making the interaction indirect. While this type of cyclic analysis adds information, interpretation may require care because of the possible involvement of more residues than the two or three that are considered. The nature of the interaction may involve small conformational changes (side chain shifts, for example, even if the S4 backbone remains stationary) as well as electrostatic interactions, and rearrangements of hydrogen bonding, in addition to proton transfers. The triad that we discuss later shows how interactions may be complex, and the triad does not exist in isolation; if a proton is transferred, it would have to proceed through more residues. Our triad is linked through one member to another triad, or the two groups could be thought of as one pentad, with the five amino acids as a single entity acting to transfer a proton.

## PROTON DELOCALIZATION AND A TRANSITION THRESHOLD

Protons are fairly light particles, even at about 1836 times the mass of an electron, and can be delocalized over a small distance, of the order of 1 Å. In addition, there can be a resonance-like proton ring formed with an acid, a base (*i.e.,* guanidinium) and two water molecules (117). A somewhat related structure has been reported by Dong et al in the influenza M2 proton channel, with four histidines and a tryptophan in a hydrogen bonded configuration, with water molecules nearby (118). These authors do not consider the possibility of a ring with resonance; they do, however, consider tautomerism; they do not give energies for each tautomeric form, but the cost for releasing a proton to water of 27 kJ mol^−1^ is the same for the two tautomeric structures. It is also possible for protons to tunnel about 1 Å, and by doing so cross a barrier that becomes available as depolarization proceeds. As mentioned earlier, Green and coworkers showed that threshold crossing could produce a gating current-voltage curve essentially identical to the experimentally determined curve if one assumes a distribution of threshold energies of width approximately k_B_T(81, 82). Proton transfer, possibly proton tunneling, would be a candidate for such a threshold crossing (119).

## COMPUTATIONAL TECHNIQUES: GENERAL CONSIDERATIONS AS TO CHOICE OF METHOD

There have been multiple attempts to calculate all or part of the ion channel, in various aspects of its structure and function. Calculations have been done for the gating mechanism, or the response to a change in voltage, as well as inactivation, selectivity, and conduction. There are essentially electrostatic physical models of ion permeation through the pore, largely proposed by Eisenberg and coworkers (120-123); these tend to use a simplified model of the channel, in which many of the specific interactions are omitted, or lumped into a smaller number of parameters; it is interesting that electrostatics can account for so much of the description of ion permeation. That said, it still will not be sufficient for the explicit description of the interactions involved in ion transport, nor is that the intended purpose of such an essentially macroscopic view. There are also much more detailed, atom level approaches. The various advantages, or difficulties, of different methods vary for the different possible systems that need to be calculated. The question has been considered in detail, and in generality, by Ahmadi, et al(124).

The most common calculation technique is molecular dynamics (MD). MD has some significant advantages: it is possible to use over 100,000 atoms, thus including lipids and hydration, and to run the simulation for times up to the microsecond range, approaching times of biological relevance. There is little doubt this will improve as computers become more powerful. Woelke et al considered the proton transfer in cytochrome c that is analogous to the channels we are discussing here(68). Examples of calculations on ion channels with particular relevance to gating include Delemotte et al, Deyawe et al, and Tronin et al (125-127). The latter does not agree with the first two. Jensen et al have carried out some extremely large simulations of gating (128, 129). Sansom and coworkers have used MD for several aspects of channel properties (130-132). While MD has been used extensively, it has significant disadvantages. One problem with MD simulations of gating is that they have not as yet been shown to return to the original state upon release of the voltage—only steered MD appears to have been tried, and that is not a proof that the system as modeled in the MD simulation is capable of returning and repeating the cycle. This is critical; If any calculation fails to show how the VSD can return to its closed state accurately multiple times, the calculation cannot be entirely valid; above, we gave 24,000 cycles per channel protein lifetime as a conservative estimate. It is not clear how the MD simulations could return correctly even once, given the large numbers of salt bridges and hydrogen bonds that must break and reform exactly correctly each cycle, in the versions we are aware of; no MD simulation to date has shown the VSD as maintaining its structure while going through even one complete gating cycle; also, some seem to require fields an order of magnitude larger than physiological to force some motion; with realistic voltages S4 does not move. Large conformational transitions are not impossible for proteins, as in enzymes with large hinge motions that do not disrupt secondary structure. Unlike most enzymes, the motions proposed in the standard models for channels are not hinge motions, involving a single bond generally moving as a rotation, or at the most two or three, but motions that require a major rearrangement of the internal structure of the protein, including the secondary structure in many versions. Conformational changes in enzymes do not often require rebuilding secondary structure. It is even harder to see how mutant channels would allow a consistent calculation, as they would have a different set of salt bridges and hydrogen bonds, which presumably would provide a different refolding problem. None of these problems arise if the backbone is stable, and only side chain motions are possible. Yet another problem is that MD calculations are unable to deal with charge transfer in most cases; as the configuration changes, and interatomic distance changes, there will be non-negligible changes (up to perhaps 0.1*e*) on local charges, causing changes in polarization that in turn lead to discrepancies from the charges used by the force field, if the force field is fixed (not polarizable). Using Drude polarization(39, 133) helps, but does not deal with much of the rest of the problems; in addition it slows the calculation so that some of the advantage of MD calculation is lost. Another consequence of charge transfer is the change in electric field and potential in the neighborhood of the transferred charge, which again cannot be accounted for in an MD simulation in which charge does not transfer. In spite of the multiple MD simulations of gating, there is still no definitive model; the MD simulations cited above show different kinds of motion.

The main alternative to MD is quantum calculations (QM). These have the advantage that they allow for charge transfer, do not require preset charges on atoms, and naturally include effects of polarizability. Both MD and QM calculations can allow exchanges of hydrogen bonds, but only QM can allow a proton to switch the atom to which it is covalently bonded, when this provides a lower energy. There are disadvantages as well for QM calculations. To begin with, there are no dynamics. Second, the number of atoms is limited; until recently, only a small section of the channel could be calculated, but as computers become more powerful, the section that can be calculated is, as of 2018, in the neighborhood of 1000 atoms with a high performance cluster, and a fair amount of patience. This is enough that the central section of the calculation will be relatively unaffected by boundary issues, which are important in small sections. QM also produces static local energy minima, corresponding to 0 K structures; one does not get a global minimum. These structures are relevant however; the most common experimental method of getting an ion channel structure has been X-ray diffraction, which is done at temperatures below 220 K, the lowest temperature phase transition for water. The X-ray structures should therefore essentially be the same as those at 0 K. Since the X-ray structures have proven to be essentially correct, the QM structures should be also. One can test for which local minima are correct, as a function of voltage, the relevant parameter in gating, by doing multiple optimizations, starting from different initial configurations, getting multiple local minima, which should include all that are relevant. For example, one may start with different states of ionization of the acids and bases (i.e., proton positions). If there is a local minimum so that the proton does not shift, the energy of that local minimum can be compared with the local minimum found in which a proton does shift. In other words, with other ionization states of the acids and bases. If all the relevant structures are determined, the lowest energy can be found for each of several voltages. Although this is time consuming, it is not so much so as to be impractical, as computers have become sufficiently powerful (134).

QM calculations can also be combined with classical molecular mechanics to produce a hybrid calculation (QM/MM) in which a central section is done as a QM calculation, and the remainder at MM level to provide a reasonable boundary region. Another possibility is to use ONIOM, which allows three different levels of calculation, including different levels of QM calculation(135). As we have discussed, there are difficulties for each method. We have already considered MD and QM. QM/MM requires that a boundary be introduced, with the bonds normally broken and sealed with H atoms. This leaves the potentials nearby somewhat inaccurate, and if the idea is to limit the QM region to simplify and speed the calculation, the QM region should not be large, so the boundary is a problem. Still this allows inclusion of many more atoms than pure QM, as the MM calculations are much faster. *Ab initio* MD, in which quantum calculations are included as the simulation progresses, is extremely resource intensive. It usually requires too short a run to show the behavior of an ion channel, and also limits the number of atoms that can be included. Sometimes calculations on relatively small to medium size systems are useful (136-139), but in general we need to make the clusters large enough that the part we are most concerned with does not have very significant boundary effects.

## SOME INTERESTING THINGS THAT WATER DOES

Proton transfer often involves water. We will show an example that does not involve water, but there are two unusual cases that may be relevant to our problem. First, it is possible to set up a system of guanidinium (thus, functionally arginine) plus a carboxylic acid, that incorporates two water molecules between the guanidinium and the acid, so as to form a ring. When the ring is complete, the energy drops sharply. On further study of the system, the behavior is most easily understandable as a resonance hybrid structure, as referred to briefly above (117). The fact that there is a small resonance structure that could stabilize a more complex system suggests care must be taken in trying to understand a structure of protein that contains water. In a sense this is not a surprise; most proteins interact with water in one way or another.

One other property of water is possibly important here: it takes at least two water molecules to get a salt bridge to ionize (140). If no water enters the center of the VSD, one should not be surprised if salt bridges are not ionized. However, as there are other polar entities nearby in some cases, it remains possible that the salt bridge may ionize, or may sometimes ionize, with the ionization state depending on external voltage and the orientation of neighboring side chains.

There is an experimental X-ray structure at a pentameric channel gate by Sauguet et al, in which water, associated with a Na^+^ ion at a channel gate, looks like a basket of five water molecules(141), plus a fairly tight second set of water molecules within the channel pore, above the ion. This turns out to be very similar to a basket of water molecules that we have calculated for a KcsA channel gate. In the Sauguet et al study the Na^+^ ion may be trapped at the channel gate by the water interacting with the protein. This appears to be closely analogous to a structure we have calculated for the gate of a KcsA channel. The water structure we have calculated is shown in Fig, 5.

**Fig. 5:**
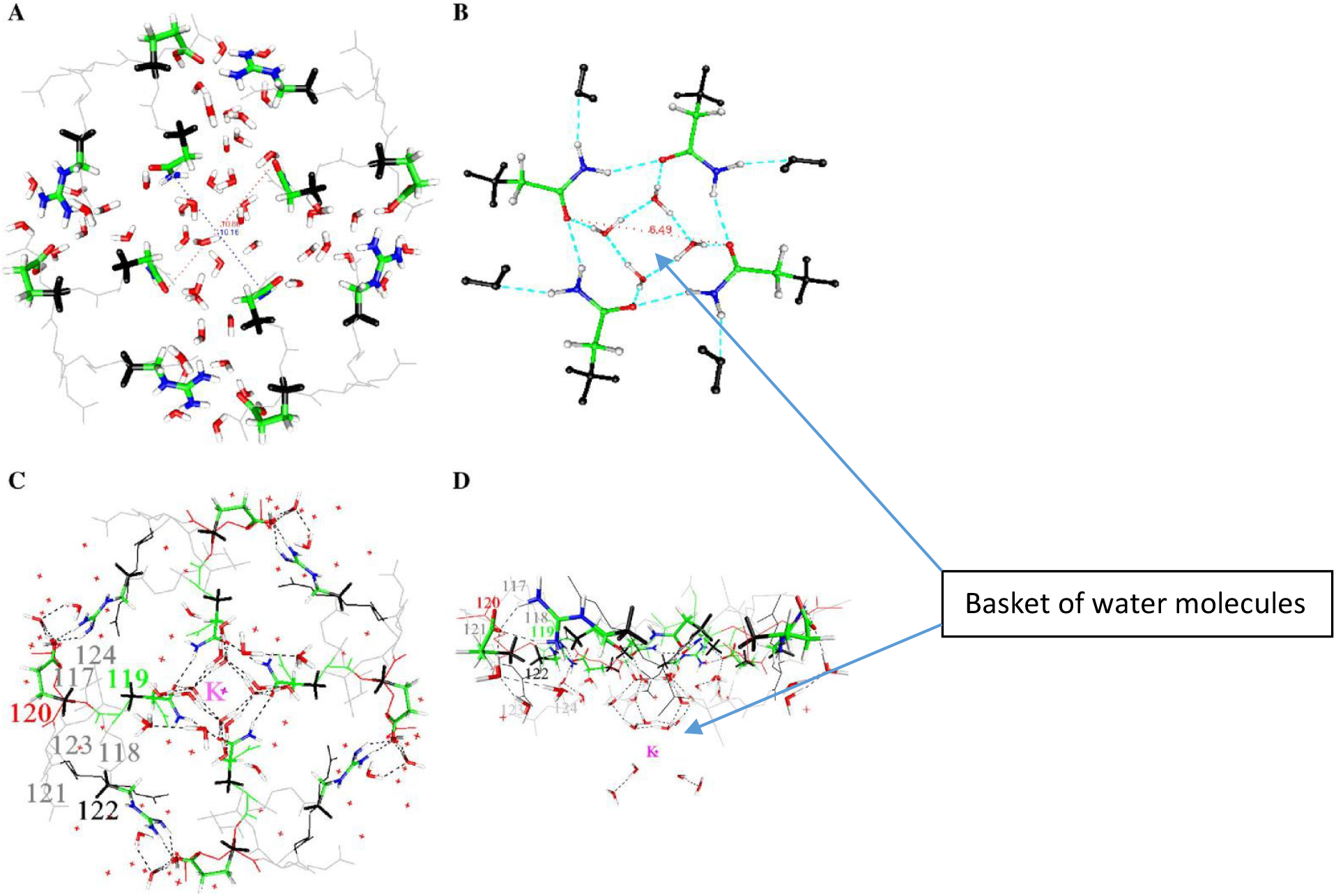
Four views of the KcsA gate: **(A**) Open (protonated) state of the KcsA channel, showing results of calculations on three amino acids from all four domains, plus water, optimized at HF/6-31G** level. Outer methyl groups are frozen. The distance between the two oxygen atoms of Q119 (a key amino acid, which may partially control the gate) is 10.9 Å on one diagonal, 10.2 Å on the other. (**B**) The closed state, Q119 only, plus the water: this calculation is done at B3LYP/6-311+G** level. Instead of 11 Å, the distance across is now only approximately 6 Å, not enough for a hydrated K^+^ ion, even if the water molecules could exchange instead of blocking the channel. **(C,D**) the uncharged (closed) state of all three amino acids (2 orientations), plus the remainder of the structure, not present in the computation. This figure is reproduced from Kariev, Znamenskiy, and Green (142).

Water also adopts a structure, based on calculations at HF/6-31G* level, that could allow it to partially(?) control a gate(142). In the KcsA channel, which is pH gated rather than voltage gated, a calculation at the gate produced the surprising structure of four water molecules shown in Fig. 5.

The calculation done so far does not show whether these molecules actually block the gate, nor whether adding one or more H^+^ nearby breaks the basket as part of opening the gate. This possibility requires further investigation from two points of view: what happens when local charge changes? Does the basket break? Second, what happens when a K^+^ ion approaches? Does the basket stop the ion from entering the channel, thus acting as part of the gate? (142). Further work to show how the water would interact with an incoming ion is required to see whether, as the oscillating gate model requires, the group can complex a K^+^ and if so, whether it can do so with smaller diameter than the maximum open diameter.

That water may be crucial is not a great surprise. Obviously it interacts with ions. In addition, it is well known to be part of a system for transmitting protons, albeit most often in the form of a water wire. In addition, it is known that D_2_O makes a difference to gating, slowing it down, at least in Na^+^ channels (143-145); it is reasonable to assume that in this respect Na^+^ and K^+^ channels are similar. The effect appears in the last step of gating (146), which agrees with what we would require in this model. The proton transport chain begins in the VSD TM section, which senses the field. Starting from the closed configuration, it responds by sending a proton cascade in the extracellular direction, pulling protons from the gate region via a path that involves water along the intracellular surface. Thus the first steps are in the transmembrane section, and the final steps involve the gate. The final steps being those affected by water in this model, it is consistent with the experimental D_2_O results. The idea that water could play a key role in gating was the first suggestion in our work (147), which considered the possibility of an electrorheological effect, in which the water became relatively rigid in the presence of a strong electric field. Whether an electrorheological effect exists at the gate is still to be determined. The field that polarizes the membrane almost certainly does not affect the intracellular water directly, but the motion of protons may change the field sufficiently at the intracellular surface that this water too responds to the field, albeit indirectly, as a consequence of the motion of protons; only protons and side chains move in direct response to the polarization/depolarization of the membrane.

Finally, the gate is likely to oscillate (see discussion with Fig. 9), and this involves changes in water density, both at the gate and in the pore cavity. If the water is tightly enough bound to the protein, there may be a single system composed of protein and water. Determining this will require further computation. In the discussion with Fig. 9 we will consider the reasons for the expectation of oscillation of the gate.

## THE ABSENCE OF TEMPERATURE DEPENDENCE IN THESE CALCULATIONS

This leaves the question of whether the results on shifts in energy with proton placement and voltage would be different if we were able to extrapolate to room temperature. We could, in principle, have different temperature dependence of energy in the different minima for each proton position. This requires that there would be different heat capacities as a function of temperature depending on the location of protons, and on the applied voltage. While the heat capacities of cases with a proton shift would not be identical, the differences should be small enough that the room temperature order of the minima, in energy, is likely to be the same.

The added energy in going from 0 K to room temperature could differ for different proton positions and voltages if the spectrum of vibrations of the protein were significantly different. However, almost all the degrees of freedom are likely to be unchanged. While the total thermal energy will correspond to well over a thousand kT (there are 976 atoms, so that there should be, in total, 2922 vibrational degrees of freedom), the *differences* should only affect a small number of these. Almost the entire system is unchanged with a local proton transfer. The differences in the energy of the minima will surely be less than the activation energy, amounting to about 20 - 25 k_B_T, for the step we have calculated in the most detail. However, even the very small number of affected degrees of freedom, say ***n***, are not going to make a difference of as much as ***n***k_B_T since these will be very similar in the two forms of the VSD (if the vibrations are fully excited, we should see a difference of 3k_B_T for each mode from the value with the mode fully unexcited). To see a difference, the modes would have to be partially excited to different degrees, which requires that they have vibration frequencies that would be in the range of k_B_T/*h* (*h* = Planck’s constant), about 6 × 10^12^ Hz at room temperature. This frequency range would apply to relatively long wave vibrations, covering much more than local modes; these are the modes which should not change with a proton shift. Higher frequency local modes differ, but these modes have energy sufficiently above k_B_T as to be not excited at room temperature. Therefore, it should be possible to trust the differences in energy—the errors must cancel in the differences, not only in the 0 K minima, but also in the room temperature results. The latter may add 1 – 2 k_B_T to the relative error, but a ±4 k_B_T error is not enough to change conclusions that are based on energy changes that are appreciably larger, by at least a factor of 4. In other words, the protein is large enough with its 2922 vibrations that it should behave somewhat as a solid, or at least a glass; therefore, many of the modes are, as in a solid, long wave vibrations covering distances over much more than local modes. The long wave vibrations will be excited by the time the temperature is raised to room temperature, but they will be the same for all local minima. The changes between cases are local, which necessarily involve short wavelengths, and with it vibrational frequencies greater than the frequency corresponding to room temperature; the local modes are generally at least 3x this frequency, so that these vibrations would contribute very little, and any alterations in local modes should not contribute much to the thermodynamics at 300 K. If the protein is analogous to a solid or glass, the local modes would be analogous to Einstein modes, and in this case, mostly with frequencies corresponding to the infrared, sufficiently above thermal energy that these modes can be ignored.

This argument applies only to the question of whether a different order of transitions would be found if the calculation were done at room temperature, not whether the absolute energy would differ from the 0 K value, which of course it would. We are saying that it would differ by essentially the same amount for each case that we studied. However, there is no question that further work is needed to fully confirm this. These considerations of degrees of freedom, and water structure vs. protein structure, are, so far, essentially qualitative arguments. A quantitative estimate of the energy difference will have to await further advances in computer capacity. Estimating free energy and other temperature dependent properties from the harmonic oscillator approximation is unlikely to be better than the qualitative estimates, as the VSD is too complex for the harmonic oscillator approximation to be any more accurate than the considerations above; mode coupling is likely to be extensive, and the spectrum of frequencies not simply Debye like, although that would be a zeroth approximation, although probably not all the way to room temperature. The modes are not a sum of independent local modes.

The temperature dependence is potentially the most serious objection to quantum calculations that give 0 K energy minima, creating one possible advantage of MD calculations. The minima are local minima; they show proton positions that are steps in the path that the protons take, and are metastable. There is no relevant global energy minimum. If the system fell into such a minimum, it is hard to see how it could go forward. There appears to be no way for the system to reach such a minimum, however; also, during gating the voltage keeps changing, so the states are necessarily transient; if they were not, there could be no proton current. We can in principle estimate the lifetime from the activation energy that it would take to get from one state to another, but we cannot yet get enough states to fix all the activation energies with sufficient accuracy, and the pre-exponential terms are also hard to estimate. The apparent activation energy of the system fits reasonably well with the results to date.

## PRESSURE

There are pressure sensitive channels (e.g., MscS, MscL) that are stretch activated. However, for the channels we are considering here, relatively little work has been done on pressure effects. More has been done on sodium than potassium channels, but in all probability the effects should be quite similar. Heinemann et al found hydrostatic pressure slowed gating, with an activation volume of 33, and then 39, Å^3^ for two cases(148). There has been more work on osmotic pressure, which might be expected to behave quite differently, as hydrostatic pressure increases the free energy of the water, while the water free energy drops in case of hyperosmolarity. Zimmerberg et al found, for potassium channels, rather large solvent inaccessible volumes (>1000 Å^3^)(149). However, the change in gating current found by Rayner et al(150) for sodium channels with osmotic pressure, while in agreement to within a factor of 2 with the solvent inaccessible volume found for potassium channels, found a limited effect on the voltage response of sodium channels. They proposed a model in which there was very weak coupling between the solvent sensitive and voltage sensitive mechanisms of gating, with coupling of the order of 0.1 k_b_T sufficient to account for their data. This does not appear compatible with the large motions suggested for S4 in standard gating models. The order of 10^3^ Å^3^ volume that becomes accessible to solvent on channel opening, from the Zimmerberg et al result, corresponds to roughly a 10 Å diameter sphere. This would be consistent with the pore, mainly cavity, volume, becoming accessible. However, there is no direct evidence on this point. The weak coupling with the voltage response is compatible with very limited change in solvent accessibility in the VSD. It is difficult to see how a major allosteric rearrangement of the VSD could fail to have a much larger effect on the solvent access; it is even more difficult to understand the weak coupling of solvent to voltage effect if the VSD were involved in making volume changes on change in osmolarity. If a large section of the VSD were to become available extracellularly on gating, a coupling of at least one hydrogen bond energy, hence >k_B_T, would be expected. The Rayner et al result is compatible with the VSD remaining largely unchanged in structure upon gating; it may be consistent with the pore becoming available.

## TOXINS

A number of toxins, e.g. particular toxins from specific snails (151-153), scorpions(154-158), and spiders (159-161), interact with voltage gated ion channels, sometimes with an extracellular loop of the VSD, or else possibly near the membrane interface. Suggestions of how these interactions with channels are relevant to gating have been proposed. We omit discussion of these in this paper; while these are a very likely useful way to obtain information as to gating mechanisms, the interpretations are not entirely obvious, and it would take a considerable amount of discussion to consider exactly how these results accord with the various possible gating models; we have not discussed conduction in detail, and some toxins interfere with conduction rather than gating. While potentially valuable, this information does not yet seem to allow a specific interpretation at the level we are considering.

## RESULTS OF RECENT QUANTUM CALCULATIONS ON THE VSD

### 1) Choice Of System For Quantum Calculations

Choosing the system size is critical to start; if too small, the result will be unrealistic, if too large there will be no result at all in a finite time. Boundaries matter. We avoid most of the difficulties of the boundary by using a large cluster (976 atoms in the first results, which we review here; an 1180 atom calculation is in progress). Using many fewer atoms (672 atoms) led to the collapse of the structure, and a clearly nonsensical result; too many of the side chains had been deleted, leaving a cavity in the center of the VSD. The surrounding residues, including the backbone atoms, collapsed into this cavity. While not done for this purpose, this also helped show that the backbone is not inherently immobile in these calculations. (It also suggests that an R➔C mutation, which leaves a volume of the order of 100 Å^3^, could be subject to collapse, so that the SCAM results must be evaluated with this in mind.) The 976 atom structure contained all the side chains that could interact with each other. There is no cavity, so no collapse, and we observe no vertical motion of the backbone of S4 either. The central region of the cluster in this case is far enough from the boundary that we can be reasonably confident that the changes calculated on transferring protons in the center are accurate. We have already discussed the fact that what we get is the energy minimum, which corresponds to 0 K. We noted this does have one advantage, in that it allows comparison with X-ray structures that are determined at temperatures below the lowest phase transition of water, and are thus presumably not very different from the 0 K structures. Cryo EM structures are of course also determined at low temperature. (An alternative, starting from snapshots of MD simulations(124), and then finding an energy minimum, seems to combine disadvantages of both methods.) By using a larger structure we minimize boundary issues.

We have carried out quantum calculations on 29 states of the 976 atom section (904 atoms of protein plus 24 molecules of water in a cleft at the extracellular end) of the VSD of K_v_1.2, where “states” refers to some combination of the field on the system and the position of a proton that can occupy positions on one of several side chains.

### 2) Results

A triad of residues, Y266, E183, R300, exchanged a proton when the membrane depolarized. The triad is very similar to a triad in bacteriorhodopsin from the 1.52Å resolution structure of Lanyi et al (rcsb pdb: 1p8H) (71). Bacteriorhodopsin is known to transmit protons. Fig. 6A compares the triad that is present in the VSD to that in bacteriorhodopsin. None of the triad of amino acids in the calculated section of the VSD was near the border of the 976 atom section, so that calculation errors from borders should be very small. It turns out that the tyrosine is capable of ionizing, transferring a proton to the NE of R300 (using the pdb notation, this is the nitrogen on the side chain before the guanidinium; the amine nitrogens on the guanidinium are labeled NH1, NH2). The proton positions calculated include the standard arginine positive, glutamate negative, tyrosine neutral configuration (which is not the low energy configuration whether polarized or depolarized), as well as exchanges of the proton that made the arginine and glutamate neutral (membrane polarized, i.e., closed conformation), or the tyrosine negative, with its protein transferred to glutamate (neutral) with arginine positive (open, depolarized). By using the NE nitrogen as a second base (or acid, when protonated), the arginine side chain becomes amphoteric. No other amino acid has a side chain with this property. Voltages in the calculations ranged from −70 mV (i.e., intracellular negative, channel presumably closed) to +70 mV (with −35 mV, 0 mV, and +35 mV also included). The energy of the crossing point at about −20 mV was about 24 k_B_T (60 kJ) above the polarized (closed) minimum. Thus the minimum conformations are: when closed (−70 mV): Y266(-), R300(+), E183(0); open (0 mV): Y266(0), R300(0), E183(0). No water is involved.

**Fig. 6A:**
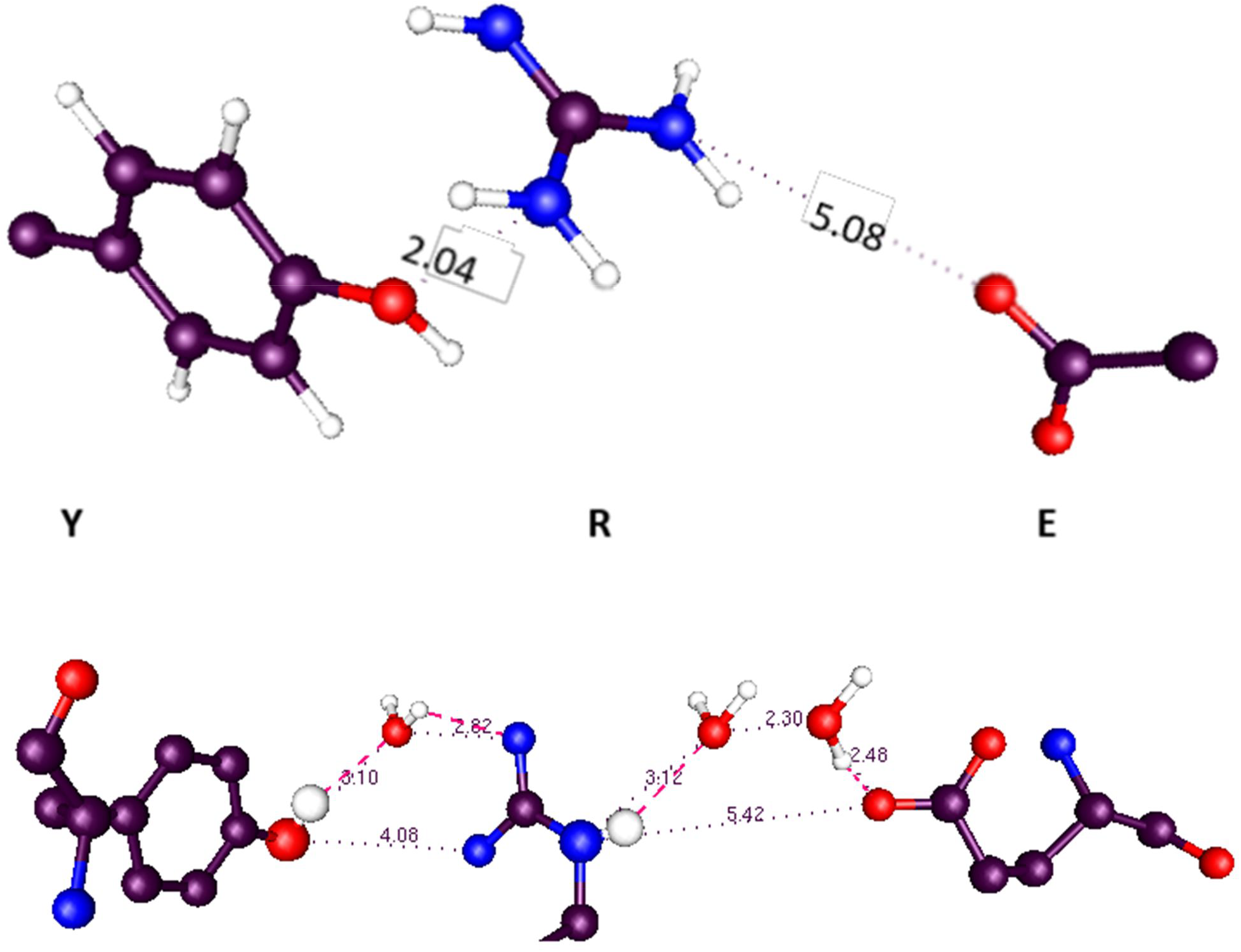
Comparison of VSD (top) and bacteriorhodopsin (bottom) triads of tyrosine, arginine, and glutamate; both are from the X-ray structures, not computation. Some interatomic distances are shown, and are very similar; e.g., the arginine nitrogen to glutamate oxygen: 5.08 (VSD) vs 5.42 (BR). Three water molecules are present in the bacteriorhodopsin structure, with one very short distance (2.30 Å) between water oxygens (oxygens are from the X-ray structure, hydrogens added), and only parts of side chains are shown. The distances are similar: especially the arginine nitrogen to glutamate, which is the same within experimental error. It is not difficult to see a proton path across the triads. Tyrosine, arginine and glutamate are on the VSD: Y415,R419,E136; on bacteriorhodopsin (pdb: 1P8H) Y57,R82,E194. This illustrates that there are several triads, as this is not the triad discussed earlier, but a different set on the path to the gate. The orientations are not identical, and the water in the bacteriorhodopsin makes up the difference.

**Fig. 6B:**
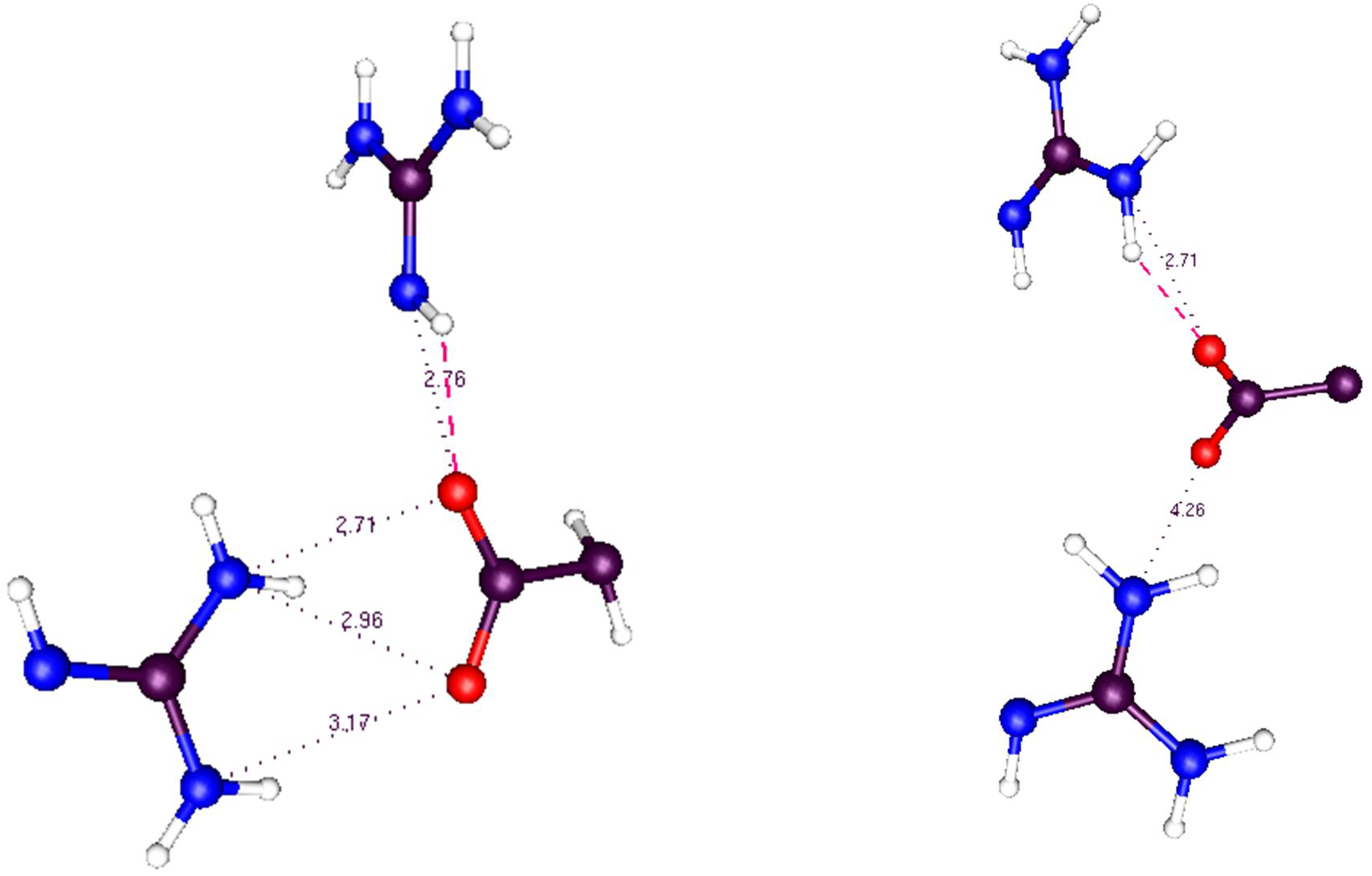
A second triad, comparing K_v_1.2 to H_v_1, this one RER/RDR instead of YRE. Left: H_v_1: R201 (top), D108,R204 (pdb:3WKV); right the VSD of the K_v_1.2 channel (3Lut); R300(top), E226, R303. H_v_1 uses the NE nitrogen of R201 (top). As in Fig. 6A the distances of the triads are very similar. The use of the arginine NE by H_v_1 is an example of the use of the amphoteric property of this side chain to transmit a proton.

The hydrophobic section of the VSD is capable of transferring a proton. There is, below this triad, a second triad that would provide a next step in the path for the proton, in going from open to closed. These are linked through the Y266, the other two members of the second triad being R303 and E226. We can propose a path all the way from this group through to the gate. The triads shown in Fig. 6 are further along in the path than the YRE triad discussed above.

In the section in the middle of the calculated region (the Y266, R300, E183 section), there are two possible pathways for the proton to go through, either via the tyrosine Y266, or the glutamate E226. The two possible proton pathways are shown in Fig. 7. The difference in energy of the states is comparable to the exchange plus correlation energy differences. For the closed configuration (nominal charges: Y266(0), R300(0), E183(0)), exchange plus correlation energy summed to 96.0 kJ above the lowest energy state (total energy was much larger), while for the lowest energy open state (Y266(-),R300(+),E183(0)), it was only 25.2 kJ. The much larger electrostatic and kinetic energy terms more than made up for the differences, but if these relatively small differences in XC terms were neglected, the states would still be placed out of order.

**Fig. 7:**
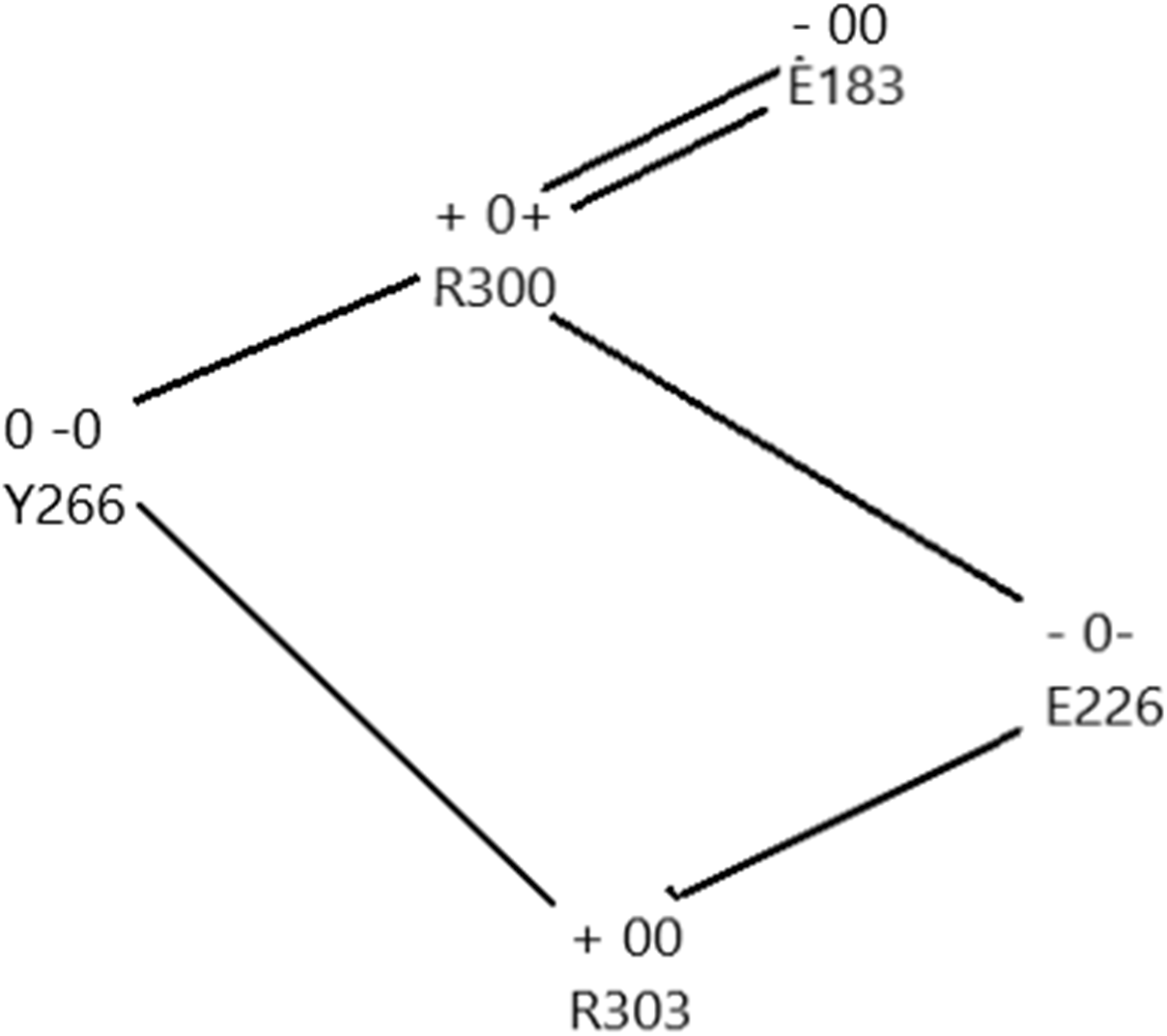
Two paths from R303 to R300, and then on to E183. The single lines represent one of the paths, the double lines mean both paths follow that step. The left symbol gives the nominal charge on that amino acid in the closed state, the right two symbols give the charge in one path each, with the left charges all for the same path, and the right charges all for the same path (e.g,, E183 – 00 means E183 is negatively charged in the closed state, and neutral in both paths to the open state). Actual charges as calculated by NBO are fractional, but of the sign shown. The proton follows the charges as shown. In the closed state, the S176 –OH is oriented so as not to form a hydrogen bond with R303 in the closed state, and oriented to form such a bond in either path of the open state.

The existence of two paths means that one path can be mutated out of existence, and the channel still functions, albeit not as well. A Y266F mutation shows just this effect. The left side path includes a tyrosine that has to ionize to make that pathway work. It appears that mutating this to a phenylalanine must interrupt this path, and make it more difficult to open the channel. The experiment has been carried out by Dr. Carlos Bassetto and Professor Francisco Bezanilla, and it showed approximately a 1.7 fold drop in slope of the gating current – voltage curve, showing that eliminating the possibility of ionizing this residue did interfere with gating, as predicted. In the standard models, one might expect, if anything, that given the slightly smaller volume of phenylalanine, a Y266F mutation might increase the probability of gating, rather than decreasing it, because it would be easier for S4 to slide up. Since the E226 path remains, the channel still functions, but not as well. This evidence supports the model we propose, and tends to refute the standard model.

There must be a gating charge, so that there must be a shift in the center of charge. For the 976 atom case, we used NBO to find the charges on all the atoms. Then, by dividing the system into 11 3 Å slices, and adding all the charges on the atoms in each slice, we determined the center of charge in the two cases of low energy (−70 mV, Y266=R300=E183=0 nominal charge, 0 mV Y266 negative, R300 positive, E183 zero nominal charge). For the −70 mV case, the center of charge was at 19.6 Å, for the 0 mV case, at 8.3 Å, for a shift of 11.3 Å. This is very nearly just what it should be, but it must be taken with caution, as the error bar on this value may be several Angstroms. The total charge on the system is always +2.

## GATING MODEL FROM THE COMPUTATION

If proton transfer provides gating current we should find a way in which intracellular protons could close the channel, while protons that have left the intracellular section would allow it to open. We note that near the gate there is a histidine (H418 in the 3Lut numbering) on the C terminal end of the pore helix S6, which could affect gating if charged. If this histidine is deleted, the channel stops functioning(162).

There are also two glutamates on T1 in the neighborhood. While none of these shows conclusively that there is a spot at which the proton chain stops, they do show that there are possible places for the protons to act on the channel gate. Further work will be required to show which of these residues acts, or which combination does. The presence of proton acceptors adds to the plausibility of the model.

The fact that T1, the intracellular section just below the membrane, is involved in gating (22, 163, 164) is itself interesting. Several suggestions as to how to integrate this into standard models have been made, although it is not obvious how to do so. T1 sits partially below the VSD and has an unusually polar interface. Also, given the geometry of the system, there is space for water molecules at the bottom of the VSD, just above T1. These waters help form a path that could conduct protons. Wang and Covarrubias (164) found that the cysteines in T1 react at very different rates with an MTS reagent; reaction depends on ionization of the cysteine, which suggests a possible reason for the difference in rates; it also suggests a role for a proton, since the ionization of the cysteine requires the transfer of a proton. T1, together with the water at the intracellular surface, appears to create a path for protons; this does not necessarily include a role for the ionization of cysteine in T1, but it does make it probable that the ionization of T1 cysteines may be important. We can hypothesize a path to (toward) the gate from the section we have already computed by examining possible hydrogen bonds that could create a path for the H^+^ to follow. Fig. 8 suggests how this would be possible.

**Fig. 8:**
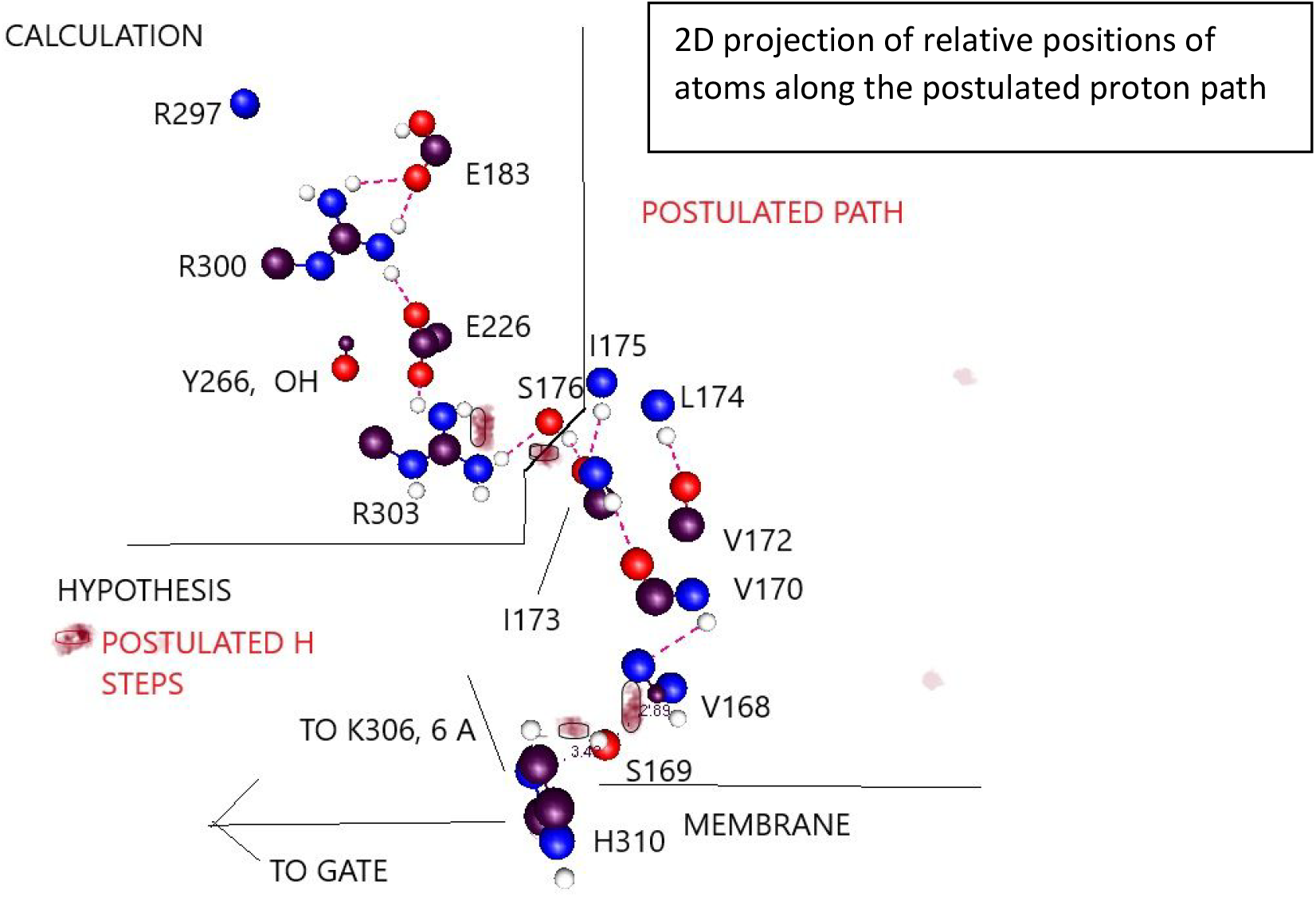
A postulated path for the protons from the calculated section in the upper left toward the gate, which would be in the lower left, off the diagram. Almost all atoms of the amino acids are omitted, but the two dimensional projection of the geometry is suggested by the diagram, which shows key atoms for each amino acid, and backbone atoms for the hydrophobic residues. Residues, all hydrophobic, from the S1 helix are shown to indicate the geometry, but there is very likely to be water in the neighborhood, between the helices. Since the path is not perfectly defined, a smudge (red) is used to indicate approximate step paths. On depolarizing the membrane, the proton would follow a path from the lower left, to the lower right, and then diagonally into the calculated region at the upper left. The S1 helix begins and ends nearest the residues at the ends of the path (R303 is near I173, and H310 is near the other end of the groups shown). It is encouraging that a very reasonable value was found. Atom colors: black=C, red=O, blue=N, white=H.

In Fig. 8, the postulated path shows only the S1 hydrophobic residues, but there are neighboring helices, and very likely water between them. Oliver and Deamer showed increased H^+^ permeability through a membrane with a group of alpha helices of hydrophobic residues(165), analogous to what we postulate here. Beyond the section that we have already calculated, we must show how a proton cascade can reach the gate, if the gate opens as a consequence of deprotonation, or closes with protonation. In Fig. 8 the path for protons that proceeds from the section we have calculated through an intracellular path to the gate includes T1, which, as noted earlier, has been shown to be involved in gating. This part of the path does involve water. The calculated section is voltage dependent, while this intracellular section is expected to have little direct voltage dependence, although it would shift protons depending on the state of the channel because of the change in protonation of the residues above it, which do respond to voltage shifts, in a sort of domino cascade. As the voltage sensitive protons shift, they pull/push the next members of the cascade.

## THE STRUCTURE OF THE GATE

Finally, when we get to the gate itself, we observe that essentially all models assume that there is an increase in the diameter of the gate to admit K^+^ ions. There is debate as to the size of the opening, or the extent of the opening compared to the closed conformation; a quite small increase in diameter would be adequate, of the order of about 4 Å(166), meaning a 2 Å displacement per VSD.

There is also a cavity in the pore in which an ion is found in the X-ray structure. An incoming potassium ion would have to interact with such an ion, which could knock *back* an incoming ion that moved beyond the entrance to the pore toward the cavity. This ion has no particular force pushing it into the gate; there is a concentration dependence of the free energy of the ion in the external solution(102) (see Fig. 3) that suggests a free energy gradient on K^+^ that might help somewhat, especially with getting the ion into the gate. We have done a limited calculation of the interaction potential that suggests that the water so substantially reduces the interaction energy between an ion at the gate and the cavity ion that, as long as the ion at the gate stays at the gate, the repulsion energy is almost zero. However, the ion then cannot advance, or displace the water or the ion in the cavity. Therefore, we proposed that the gate would have to oscillate, complexing and holding the incoming ion, which then cannot be knocked back, so that the ion in the cavity would proceed to the bottom state of the selectivity filter, given a little time, once the bottom state of the selectivity filter becomes available. Once the cavity cleared, the ion at the gate could move into the cavity, not so much having pushed the previous ion forward, as having simply allowed it to move, and then replaced it. The cycle is indicated in Fig. 9, reproduced from (40). This implies that the gate can complex K^+^ well enough to force the ion in the cavity to move up, or at least allow it to move up. The complex must not be so tight that, once the cavity becomes available, it cannot release the K^+^; the cavity location is then of lower energy than the solution, so that the ion moves into the cavity rather than back to solution, and the current continues. Here it is interesting to compare the KcsA closed structure to the K_v_1.2 open structure, for both of which X-ray structures exist. Kariev and Green noted that the gate opening was about a 3 Å increase in the radius of the intracellular gate (167).

**Fig. 9:**
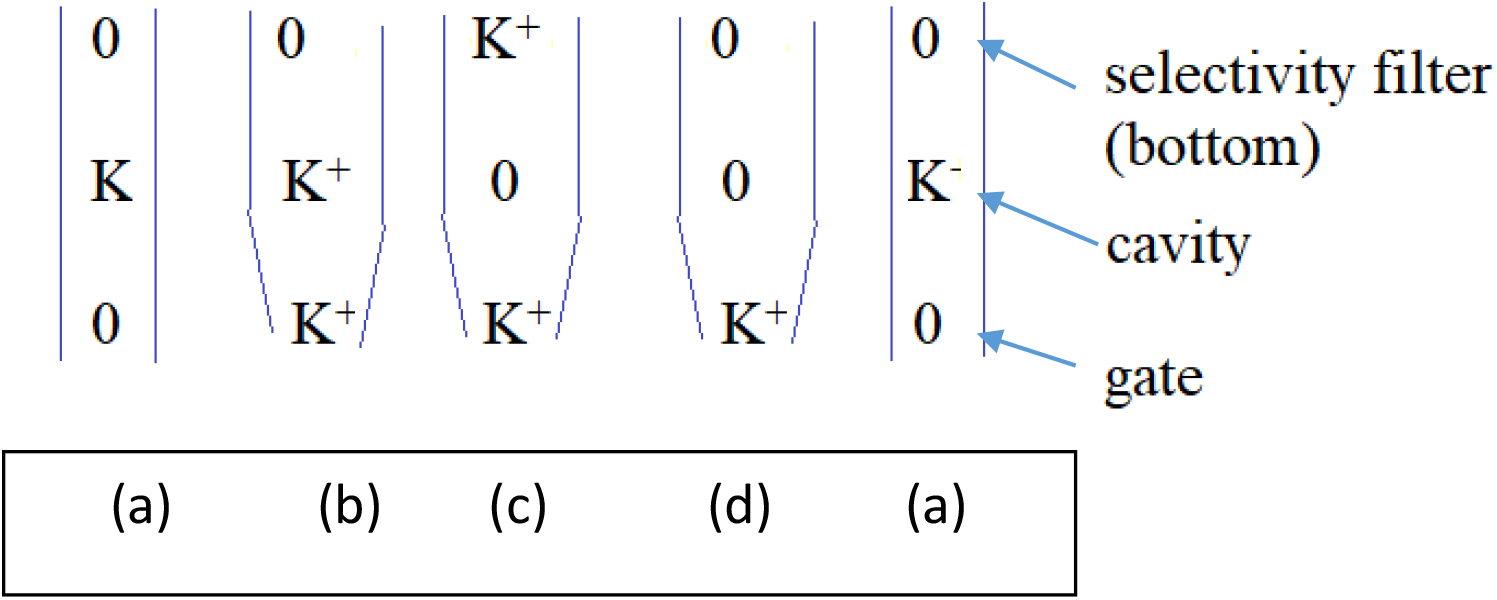
Cycle of states at the oscillating gate. In going from (a) to (b), the gate oscillates to hold the ion, which must remain until the ion in the cavity (which does show in the X-ray structure) moves up to the selectivity filter (top position, (c)). This ion can then move up further within the selectivity filter (d), leaving space for the ion at the gate to move into the cavity, returning to state (a), where the gate can again relax, in preparation for the next ion. Based on calculations the contraction part of the oscillation cycle appears to be an increased density of water at the gate, rather than a change in protein conformation or gate diameter. Adapted from ref.(40)

Then the 15 Å N-N distance (between nitrogens of two prolines from opposite VSDs in K_v_1.2) would be consistent with the entrance of a hydrated K^+^, with the ion not yet complexed by protein. A calculation of the pore region, with a total of 50 water molecules in a total of 870 atoms showed that there was no difference in the N-N distance with a K^+^ at the gate, but about three more water molecules clustered there, for a total of thirteen, compared to about ten when the ion was just below the selectivity filter. Apparently water is responsible for the oscillating gate effect. An ion can be tightly held by the increase in the number of water molecules that are both hydrogen bonded to the protein and linked to the ion. Moving the ion relaxes the water conformation, allowing another ion to approach.

Both KcsA and K_v_1.2 have linear ln[K^+^] vs ln σ curves, which should correspond to a barrier, as we discussed earlier (Fig. 3). If we compare the need for a barrier to the steps for an oscillating gate, the most obvious step to have the barrier is a➔b in Fig. 9. Here an ion from solution must enter the gate, bring the gate, with its water, to the position that complexes the ion, and rearrange the water in the cavity. Thus the model is consistent with the data; since the free energy gradient should affect an ion at the gate, but not further up the pore, the concentration dependence seems to require that that the barrier demonstrated by Fig. 3 be at the intracellular end of the pore.

### Summary on VSD calculation

The energy, atomic charges, and bond order in the VSD calculation depended on both the proton position and the applied field. The net result was that the energy for the closed (−70 mV) state is lower with arginine, glutamate, and tyrosine all neutral, while the open state has a proton transferred from tyrosine to arginine (tyrosine negative, arginine positive, glutamate neutral). This calculation is being extended to the remainder of the proton path; a plausible path has been postulated.

## SPECIFIC COMPUTATIONAL METHODS

In the QM calculations reported here, we have obtained the structures of the energy minima by optimization at HF/6-31G* level. This includes exchange but not correlation energy, in addition to all those terms that have classical analogues (kinetic energy, electron-nuclear interaction, and electron-electron and nuclear-nuclear repulsion). However, the omitted correlation energy, while the smallest term, is still not negligible in determining the differences in energy of different states. Therefore, the final energy calculations, and all calculations of properties, used B3LYP/6-31G** for necessary accuracy; as a DFT method, it includes the correlation energy. This level of calculations included NBO determination of bond strengths and atomic charges. These charges were used for calculation of the local potentials, and helped determine the position of the center of charge in each relevant state. The center of charge calculations used a program written in the group.

## SUMMARY

We have considered the evidence concerning voltage gating, principally of the K_v_1.2 channel, but also considering analogous systems. We have reinterpreted results of multiple experiments and our own calculations to show that they do not provide convincing evidence of significant S4 motion as in the standard model, but can be understood as providing evidence for the motion of protons as gating current.

Major points include:

1. There is a standard model in which gating current is provided by the motion perpendicular to the membrane of the S4 TM segment of the VSD, supported principally by:

a. SCAM studies of this and related channels
b. MD simulations
c. FRET and LRET studies
2. We have offered an alternative interpretation in which the gating current is provided by proton motion. This is supported by:

a. Quantum calculations showing how a proton can make a local transfer in the K_v_1.2 channel VSD; this depends in part on the fact that the side chain of arginine is amphoteric, unlike the side chains of all other amino acids. The amphoteric nature of the arginine side chain is unique among natural amino acids.
b. Extending these calculations to show that charge transfer is consistent with known gating charge.
c. Providing a hypothesis that shows much of the remainder of a path that a proton could follow from gate to the extracellular surface of the membrane.
d. Considering several experiments that are typically ignored or interpreted in contradictory fashion in order to fit them to the standard model, when more natural interpretations are consistent with proton transport.
e. Consideration of analogous systems, such as H_v_1, cytochrome c, bacteriorhodopsin, and the flu M channel that are known to transmit protons, It is of particular interest that all of these, like the VSD of the K_v_ channel, have the same tyrosine-arginine-glutamate arrangement in the apparent proton path, as well as other similarities (e.g., an RER/RDR triad) that would create a proton path. It would be of interest to investigate other proteins that transmit protons to determine whether this triad is a recurring motif in proton transport in proteins.
f. The K_v_1.2 channel at least has an increase in water density at the gate when an ion is present. The KcsA channel has an interesting water structure at the gate. Although water is not involved in the triad proton transfers, it is critical to the behavior of the channel.
g. The center of charge shifts very nearly just as much as one would expect; this result must have a fairly large error bar attached, but it is almost exactly what should be expected.

Taken together we have considered as much as possible of the relevant evidence, and found that the evidence supports proton transport as the source of gating current better than it supports the standard models.

## ACKNOWLEDGEMENTS

For our calculations: we thank the Center for Functional Nanomaterials at Brookhaven National Laboratory, a DoE facility, and the CUNY High Performance Computer Center, supported by NSF, for computer time. We thank Dr. Carlos Bassetto and Professor Francisco Bezanilla for carrying out the experiment on the Y266F mutant.

